# Phosphodiesterase 1A physically interacts with YTHDF2 and reinforces the progression of non-small cell lung cancer

**DOI:** 10.1101/2024.05.21.595220

**Authors:** Chong Zhang, Zuoyan Zhang, Yueyi Wu, Yuchen Wu, Jing Cheng, Kaizhi Luo, Zhidi Li, Manman Zhang, Jian Wang, Xuesen Zhang, Yangling Li

**Affiliations:** Department of oncology, Shangyu People’s Hospital of Shaoxing, Shaoxing, Zhejiang, China, 312300; Department of Pharmacy, School of Medicine, Hangzhou City University, Hangzhou, Zhejiang, China, 310015; Department of Pharmacy, The Affiliated Hospital of Northwest University · XI’AN NO.3 Hospital, Shanxi, China, 710038; Department of Pharmacy, Ningbo First Hospital, Ningbo, China, 315010; Department of Clinical Medicine, The First School of Medicine, Wenzhou Medical University, Wenzhou, Zhejiang, 325035; College of Pharmaceutical Sciences, Zhejiang University, Hangzhou, Zhejiang, China, 310058; Department of Pharmacy, Zhejiang University of Technology, Hangzhou, Zhejiang, China, 310027; Hangzhou Lin’an Traditional Chinese Medicine Hospital, Affiliated Hospital, Hangzhou City University, Hangzhou, 311300, China; Department of Clinical Pharmacology, Key Laboratory of Clinical Cancer Pharmacology and Toxicology Research of Zhejiang Province, Affiliated Hangzhou First People’s Hospital, School of Medicine, Westlake University, Hangzhou, China, 310006

**Keywords:** Key word: PDE1A, metastasis, YTHDF2, NSCLC, STAT3

## Abstract

Non-small cell lung cancer (NSCLC) is the most common subtype of lung cancer, and the prognosis is poor due to distant metastasis. Thus, there is an urgent need to discover novel therapeutic targets and strategies to overcome metastasis. A series of in vitro and in vivo phenotype experiments were performed to investigate the role of PDE1A in NSCLC. The RIP assay, mRNA stability assay and LC-MS/MS were performed to investigate the molecular mechanisms of PDE1A in NSCLC progression. Phosphodiesterase 1A (PDE1A) has been shown to promote metastasis and EMT progression of NSCLC. In addition, NSCLC cells overexpressing PDE1A promoted angiogenesis by regulating exosome release. IL-6/JAK/STAT3 signaling pathway was highly enriched in PDE1A-coexpresssed genes, and PDE1A promoted NSCLC metastasis by activating the STAT3 pathway. GO enrichment analysis of PDE1A-interacting genes showed that PDE1A might interact with YTHDF2 and participate in m6A-containing RNA binding. The binding between PDE1A and YTHDF2 was verified, and PDE1A regulated the STAT3 pathway by interacting with YTHDF2. The mechanism of YTHDF2/PDE1A complex in regulating STAT3 pathway was predicted by overlapping YTHDF2-interacting-RNAs, and genes coexpressed with YTHDF2 and STAT3. The interactions between YTHDF2 and target mRNAs were predicted, and there were three predicted targets of YTHDF2 with high scores: NRF2, SOCS2, and MET. Indeed, PDE1A interacted with YTHDF2, destabilized SOCS2, and activated STAT3 pathway. The data not only uncovers a novel PDE1A/YTHDF2/STAT3 pathway in NSCLC progression but also provides therapeutic strategies for treating NSCLC patients with metastasis.

## Introduction

Lung cancer is one of the most frequently diagnosed cancers and the leading cause of cancer-related mortality worldwide ^[1]^. Non-small cell lung cancer (NSCLC) represents approximately 85% of lung cancers and is the most common subtype of lung cancer ^[2]^. Despite rapid advances in the clinical treatment of NSCLC in recent years, the prognosis of NSCLC patients is still poor due to recurrence and distant metastasis ^[3]^. However, the mechanism of NSCLC metastasis is still poorly understood.

Phosphodiesterases (PDEs) are a class of enzymes that hydrolyse cyclic adenosine monophosphate (cAMP) and cyclic guanosine monophosphate (cGMP), reducing the signaling of these important intracellular second messengers ^[4]^. PDEs consist of 11 family members, and each family member contains multiple subtypes ^[5]^. PDEs are being pursued as therapeutic targets in multiple diseases, including the cardiovascular system, metabolism, pulmonary system, nervous system, immunity, and cancers ^[6]^. For example, PDE3 inhibitors are used to treat heart failure and peripheral artery disease, and PDE4 inhibitors are approved to treat inflammatory diseases ^[7]^. Multiple studies have shown that PDEs, such as PDE4 and PDE5, play a vital role in the progression of tumors and are regarded as potential therapeutic targets for cancer treatment ^[7, 8]^. The PDE1 family member has three subtypes, including PDE1A, PDE1B and PDE1C, with different affinities for cAMP and cGMP ^[9]^. The effect and mechanism of PDE1 in regulating cancer progression remain elusive.

N^6^-methyladenosine (m6A) is the most abundant RNA modification, and the process of m6A modification is reversible: m6A is installed by “writers”, removed by “erasers”, and recognized by “readers” ^[10]^. YT521-B homology domain family member 2 (YTHDF2), belonging to the YTH domain protein family, has been validated as m6A “reader” and regulates the stability of messenger RNAs (mRNAs) ^[11]^. YTHDF2 promotes the progression of lung adenocarcinoma by recognizing m6A modification and influencing mRNA fate ^[12]^. Furthermore, YTHDF2 orchestrates the reprogramming of tumor-associated macrophages in the tumor microenvironment (TME), and YTHDF2 is an effective target to enhance cancer immunotherapy ^[13]^. The findings indicate that PDE1A is a promoter of NSCLC metastasis through its interaction with YTHDF2.

## Results

### PDE1A overexpression predicts a poor prognosis in lung cancer patients

First, Immunohistochemistry analysis revealed that PDE1A expression was significantly higher in lung cancer tissues compared to normal lung tissues (Figure 1A and Figure S1) ^[15, 16]^. As shown in Figure 1B, overexpression of PDE1A was also observed in three NSCLC cell lines compared with normal human lung fibroblasts (HELF cells). Additionally, the overexpression of PDE1A was also observed in lung cancer from high-risk patients compared with low-risk patients (p<0.0001, Figure 1C), and lung cancer patients in the high-risk group had shorter survival times than those in the low-risk group (Figure 1D) ^[17, 18]^. Furthermore, lung cancer patients with high levels of PDE1A in their tumors had shorter overall survival than those with low PDE1A expression, indicating that PDE1A overexpression was correlated with a poor prognosis in lung cancer patients (Figure 1E) ^[19, 20]^. Thus, PDE1A might be a novel prognostic predictor in lung cancer treatment and contribute to lung cancer progression.

**Figure 1.**
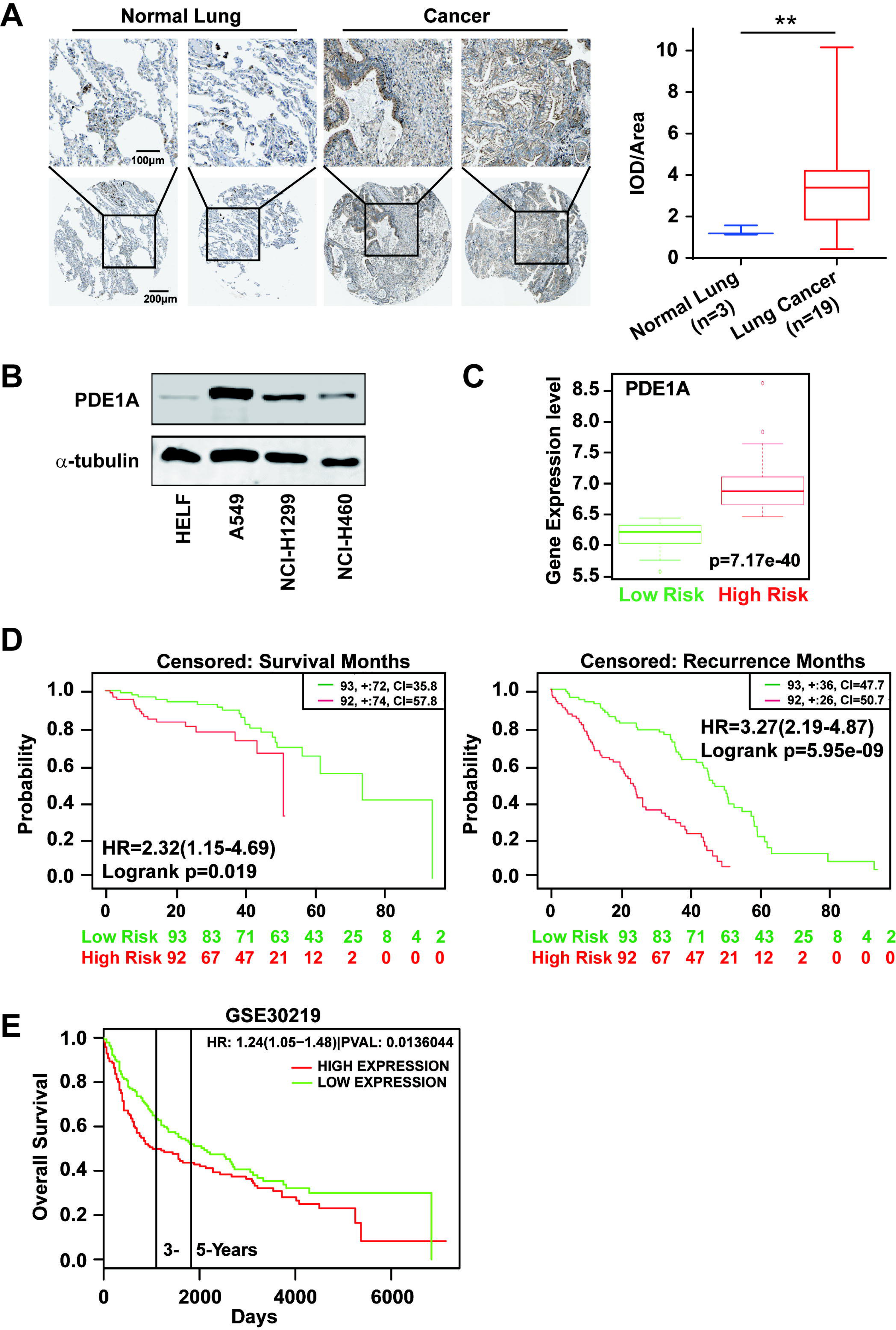
High expression of PDE1A predicts a poor prognosis of lung cancer patients. (A) The expression of PDE1A was detected in NSCLC and normal lung tissue. The data were obtained from The Human Protein Atlas (https://www.proteinatlas.org). IOD/area means Integral optical density/area. T-test was used to compare the difference between lung cancer and normal lung groups. (B) The expression of PDE1A in HELF, A549, NCI-H1299, and NCI-H460 cells was detected by western blot. (C and D) Box plot analysis of the PDE1A mRNA levels in clinical lung cancer tissue samples. The data were collected and statistical analyzed from SurvExpress (http://bioinformatica.mty.itesm.mx:8080/Biomatec/SurvivaX.jsp). Gene: PDE1A; Access database Numbers: Chitale Lung (n=185); Censored: Recurrence Months or Survival months. (E) The data was collected from PROGgene V2 Prognostic Database (http://www.progtools.net/gene/index.php). Survival analysis is done using backend R script which employs R library ‘survival’ to perform Cox proportional hazards analysis (function ‘coxph’) and to plot prognostic plots (function ‘survfit’). TSingle user input genes: PDE1A; Cancer type: LUNG; Survival measure: Death; Bifurcate gene expression at: Median; GSE30219-Off-context gene expression in lung cancer identifies a group of metastatic-prone tumors.

### PDE1A promotes the metastasis and EMT of NSCLC cells both *in vitro* and *in vivo*

To investigate the biological function of PDE1A in lung cancer development, gene set enrichment analysis (GSEA) and overrepresentation enrichment analysis (ORA) were performed to analyze the biological process of PDE1A in NSCLC using LinkedOmics ^[21]^. As shown in Figure S2A, PDE1A might be involved in the adhesion, migration, and motility of NSCLC cells, which are critical parameters in the metastatic dissemination of cancer cells. Tumor angiogenesis, the recruitment of new blood vessels, enables a pre-existing tumor to grow and metastasize ^[22]^. PDE1A might also participate in mesenchyme development, angiogenesis, vasculature development, cellular response to VEGF stimulus, blood vessel morphogenesis and development (Figure S2A-C). Thus, it was hypothesized that PDE1A may enhance the metastatic potential of NSCLC cells..

First, PDE1A silencing did not cause a significant decrease in the proliferation of NSCLC cells relative to that in the control siRNA group (Figure S3A). Moreover, PDE1A overexpression had no significant effect on the proliferation of NSCLC cells (Figure S3B). As bioinformatics analysis demonstrated that PDE1A might promote the metastatic potential of NSCLC cells, wound healing and transwell assays were used to investigate the migration and invasion capacity of PDE1 family members. Knockdown of PDE1 family members suppressed the migratory ability of NCI-H1299 cells, and siPDE1A exerted a stronger suppression effect on the migration of NCI-H1299 cells than siPDE1B and siPDE1C transfection (Figure S4A). Meanwhile, siPDE1A resulted in more profound suppression of EMT progression in NCI-H1299 cells than siPDE1B and siPDE1C transfection (Figure S4B). Thus, PDE1 family members, particularly PDE1A, might be involved in the metastatic behavior of NSCLC cells.

As shown in Figure 2A and 2B, suppression of PDE1A markedly reduced the migratory and invasive capacity of NSCLC cell lines, and the wound healing assay also showed that NSCLC cells with PDE1A knockdown had a slower wound closure rate than control siRNA-transfected cells (Figure 2C and 2D). Furthermore, PDE1A knockdown increased E-cadherin expression and reduced N-cadherin expression, indicating that PDE1A silencing suppressed EMT progression in NSCLC cells (Figure 2E). Meanwhile, the PDE1 inhibitor vinpocetine significantly suppressed the migration and EMT of NSCLC cells (Figure 2F - H). To determine the effects of PDE1A on NSCLC cell migration and invasion *in vivo*, nude mouse models were established using PDE1A-shRNA- and control-shRNA-treated NCI-H1299 cells. As shown in Figure 2I and Figure S4C-D, the number of pulmonary metastatic nodules was decreased in the PDE1A-shRNA group compared with the control-shRNA group in nude mice.

**Figure 2:**
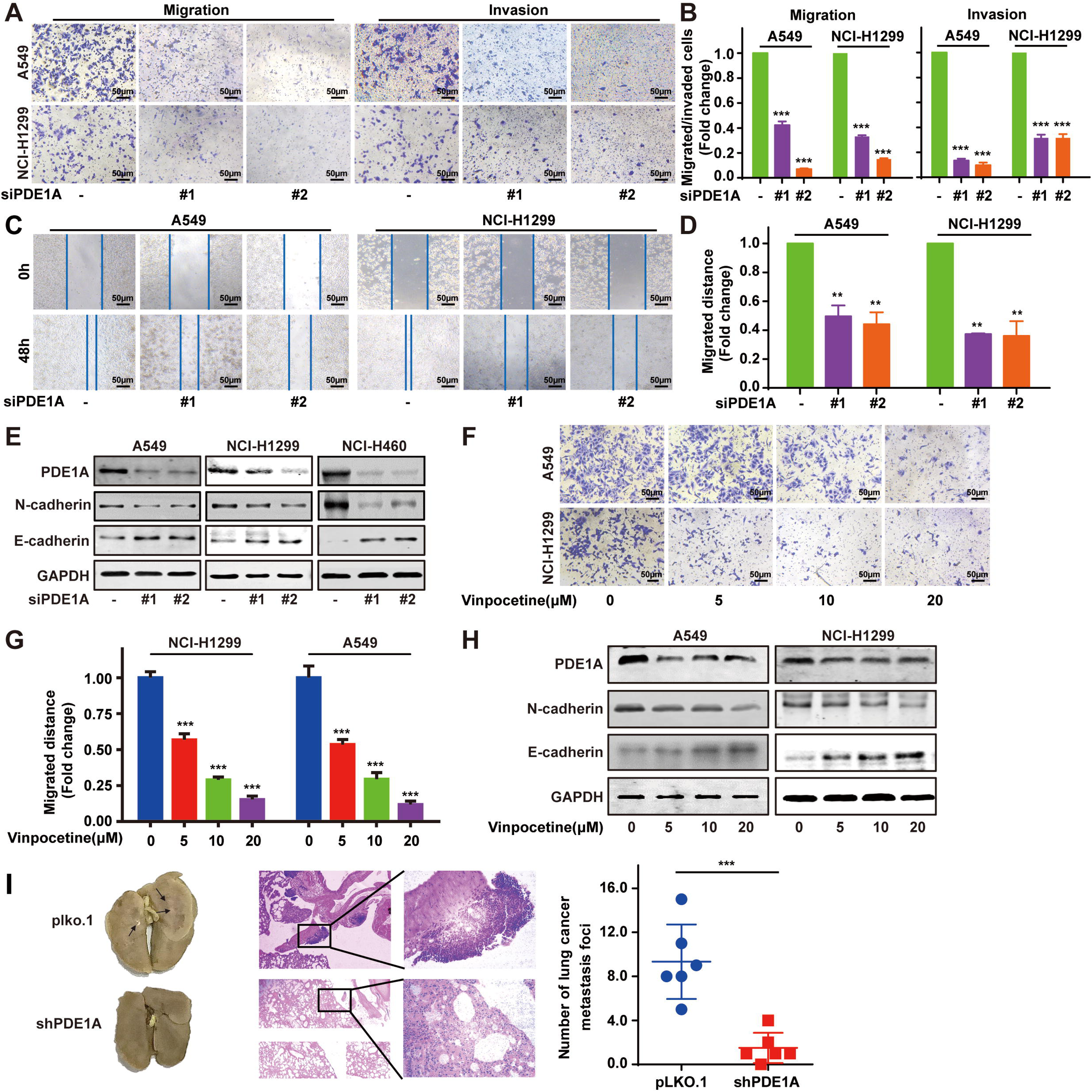
PDE1A knockdown suppresses the metastasis of NSCLC cells. (A-B) NSCLC cells were transfected with control siRNA and PDE1A siRNA for 24 h, cells were transferred to Transwell chambers without (A) or with (B) a Matrigel coating on the insert membrane, and the cell migrative and invasive abilities were determined, respectively. (C-D) NSCLC cells were transfected with control siRNA and PDE1A siRNA for 24 h, and the wound healing assay was established in NSCLC cells. (E) NSCLC cells were transfected with control siRNA and PDE1A siRNA for 48 h, and the expression of indicated proteins were detected. (F-G) NSCLC cells were treated with DMSO or vinpocetine (5, 10, 20 µM) for 24 h, and the migrative ability of treated NSCLC cells was determined using Transwell assay for 24 h. (H) NSCLC cells were treated with DMSO or vinpocetine (5, 10, 20 µM) for 24 h, and the expression of indicated proteins was determined. (I) The pulmonary metastatic nodules were stained using H&E staining and counted in nude mice harboring NCI-H1299 cells transfected with PDE1A shRNA and control shRNA.

In contrast, PDE1A overexpression significantly enhanced the migratory and invasive capacities of NSCLC cells (Figure 3A and 3B). In addition, NSCLC cells with high PDE1A expression had a higher wound closure rate than those transfected with empty vector (Figure 3C). Meanwhile, PDE1A overexpression decreased E-cadherin expression and elevated N-cadherin expression, indicating that PDE1A promoted EMT progression of NSCLC cells (Figure 3D). To compare the expression of PDE1A in NSCLC cells with high and low invasive potential, a series of NSCLC cell lines with varying invasive capabilities were established through repeated selection using the transwell chamber assay (Figure 3E, Figure S5A and S5B). The protein and mRNA levels of PDE1A were higher in highly invasive NSCLC cells than in NSCLC cells with low invasive potential (Figure 3F and 3G). In an *in vivo* nude mouse experiment, NSCLC cells overexpressing PDE1A produced more pulmonary metastatic nodules than the parental NSCLC cells (Figure 3H and Figure S5C-D).

**Figure 3:**
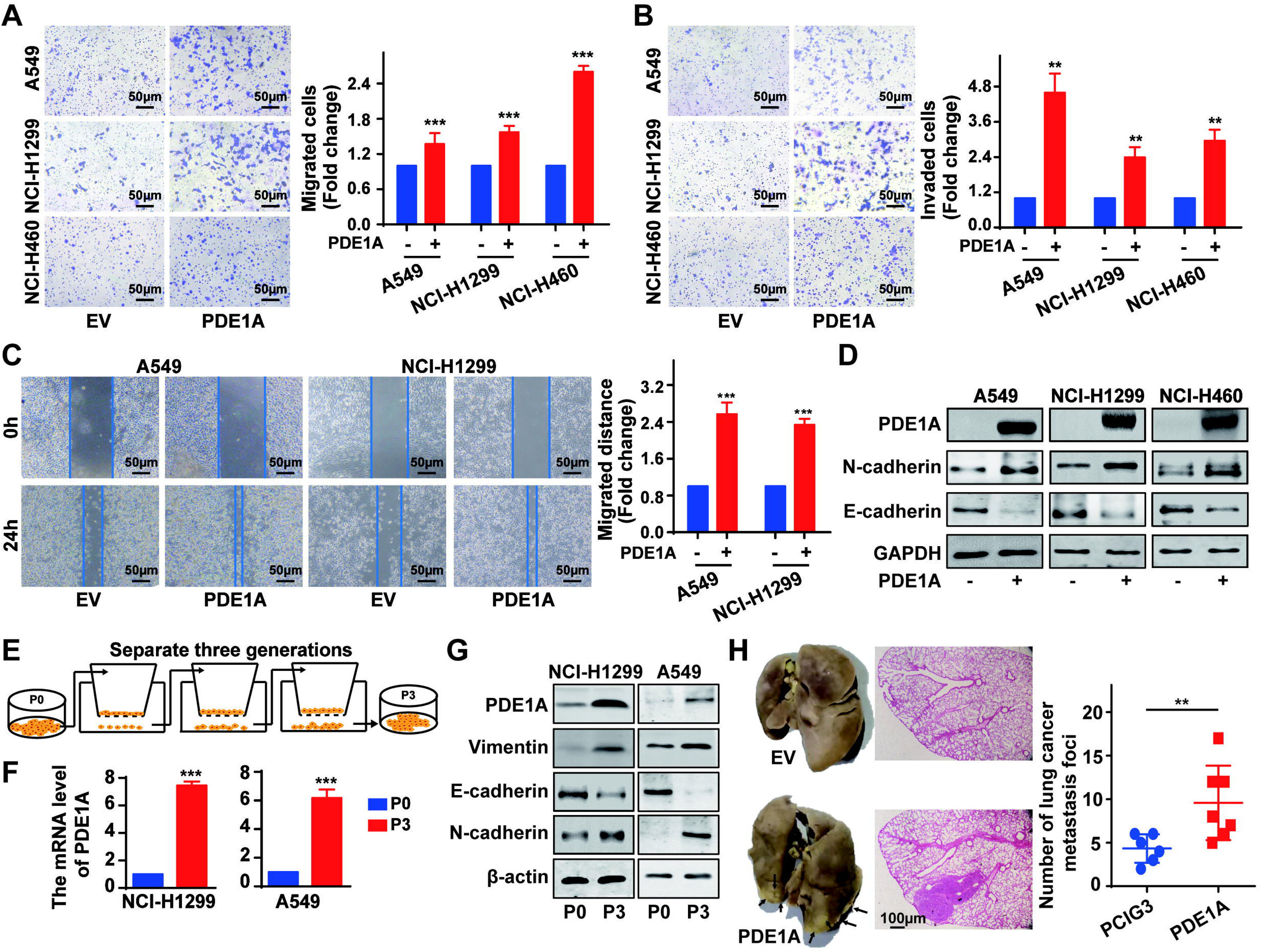
PDE1A promotes the metastasis and EMT of NSCLC cells. (A-B) NSCLC cells were transfected with PDE1A plasmid and empty vector for 24 h, cells were transferred to Transwell chambers without (A) or with (B) a Matrigel coating on the insert membrane, and the cell migrative and invasive abilities were determined, respectively. (C) NSCLC cells were transfected with PDE1A plasmid and empty vector for 24 h, and the wound-healing assay was established in NSCLC cells. (D) NSCLC cells were transfected with PDE1A plasmid and empty vector for 48 h, and the expression of indicated proteins were detected. (E) The highly invasive NSCLC cells were separated using transwell chamber assay, and P3 cells were obtained from P0 cells after three generations. (F-G) The mRNA (F) and protein (G) levels of indicated genes were determined in P3 and P0 NSCLC cells. (H) The pulmonary metastatic nodules were stained using H&E and Bouin’s solution and counted in nude mice harboring NCI-H1299 cells transfected with PDE1A plasmid and empty vector.

### NSCLC cells overexpressing PDE1A promote angiogenesis in the TME

GSEA demonstrated that PDE1A expression was positively correlated with angiogenesis in lung cancer (Figure 4A). To mimic the tumor microenvironment, a coculture system of NSCLC cells and vascular endothelial cells was established (Figure 4B). NSCLC cells overexpressing PDE1A promoted the migration of HUVECs, and NSCLC cells with low levels of PDE1A suppressed the migration of HUVECs (Figure 4C and 4D). Next, NSCLC cells were treated with GW4869 to reduce exosome release and found that GW4869 suppressed the enhancement of the migratory ability of HUVECs induced by NSCLC cells overexpressing PDE1A (Figure 4E). Meanwhile, compared with negative control, shPDE1A significantly suppressed tumor angiogenesis of NSCLC *in vivo* (Figure 4F). Thus, NSCLC cells overexpressing PDE1A promote angiogenesis in the TME.

**Figure 4:**
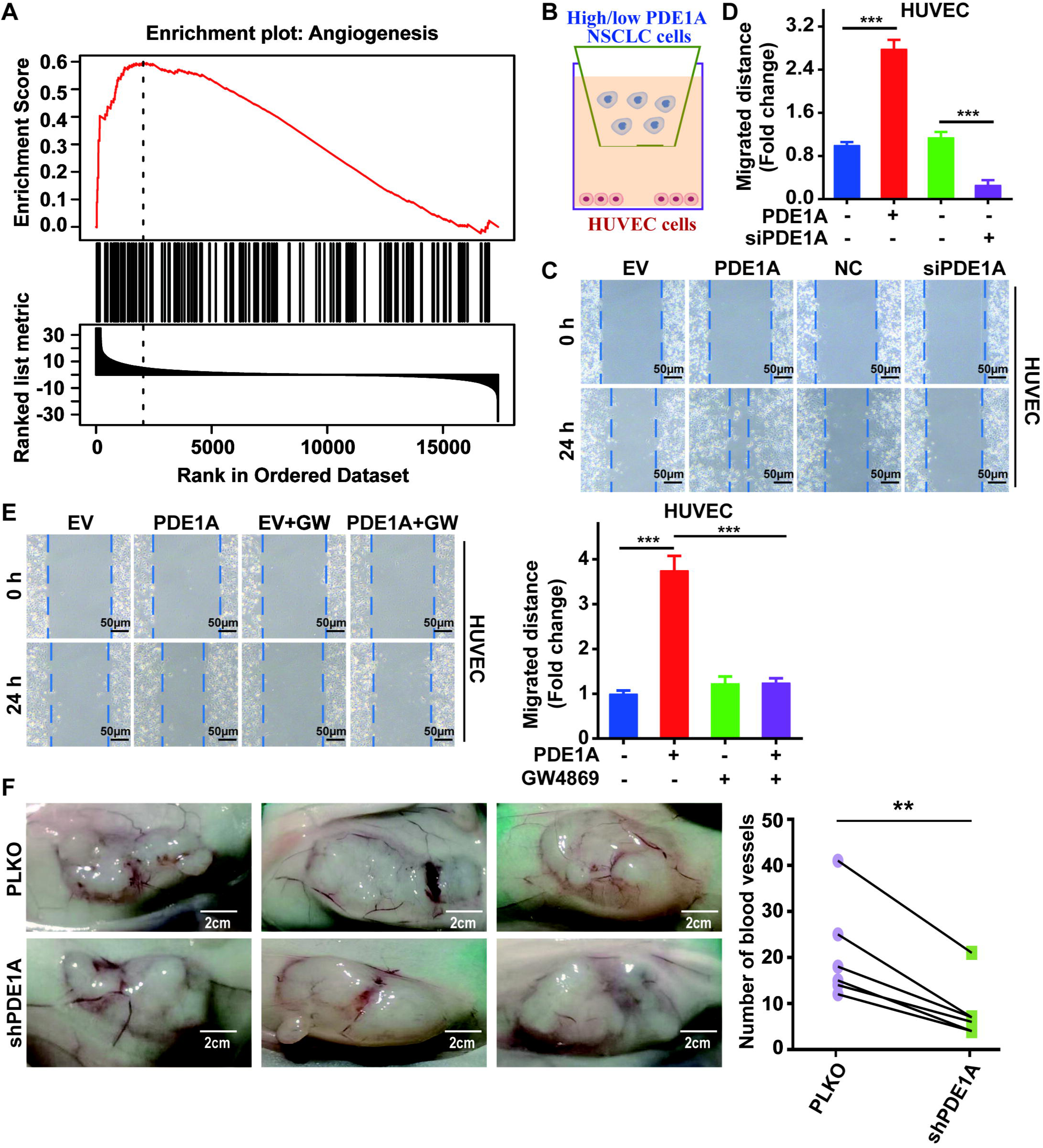
NSCLC cells overexpressing PDE1A promote angiogenesis in the TME. (A) Data were collected from LinkedOmics. Statistical tests in LinkFinder include Pearson’s correlation coefficient, Spearman’s rank correlation, Student’s t-test, Wilcoxon test, Analysis of Variance, Kruskal– Wallis analysis, Fisher’s exact test, Chi-Squared test, Jonckheere’s trend test and Cox’s regression analysis. Multiple-test correction is performed using the Benjamini and Hochberg method to generate the False Discovery Rate. (B-D) NSCLC cells were transfected with empty vector / PDE1A overexpressing plasmid or control siRNA / siPDE1A for 48 h, and then NSCLC cells were placed on the upper panel of transwell with 0.4 μm insert, HUVECs were placed on the lower panel of transwell, and wound healing assay was performed to determine the migrative abilities of HUVECs. (E) NSCLC cells with PDE1A overexpression were treated with 10 µM GW4869, and wound healing assay was performed to determine the migrative abilities of HUVECs. (F) NSCLC cells were transfected with empty vector or shPDE1A, then cells were transplanted into nude mice via subcutaneous injection, the blood vessels were counted after 60 days.

### PDE1A promotes the metastasis of NSCLC cells via the STAT3 signaling pathway

Then, the dependence of PDE1A-enhanced metastasis on cAMP metabolic activity was investigated.. As shown in Figure S6A, the cAMP inhibitor SQ22536 failed to rescue the migrative ability suppressed by siPDE1A in NSCLC cells, indicating that the basic molecular function might not be involved in the metastasis of PDE1A. To better explore the mechanism of PDE1A in NSCLC progression, bioinformatic analysis of PDE1A coexpressed genes were performed, which revealed that PDE1A might be involved in the JAK/STAT3, Hedgehog, and TGF-β pathways in NSCLC (Figure 5A).

**Figure 5:**
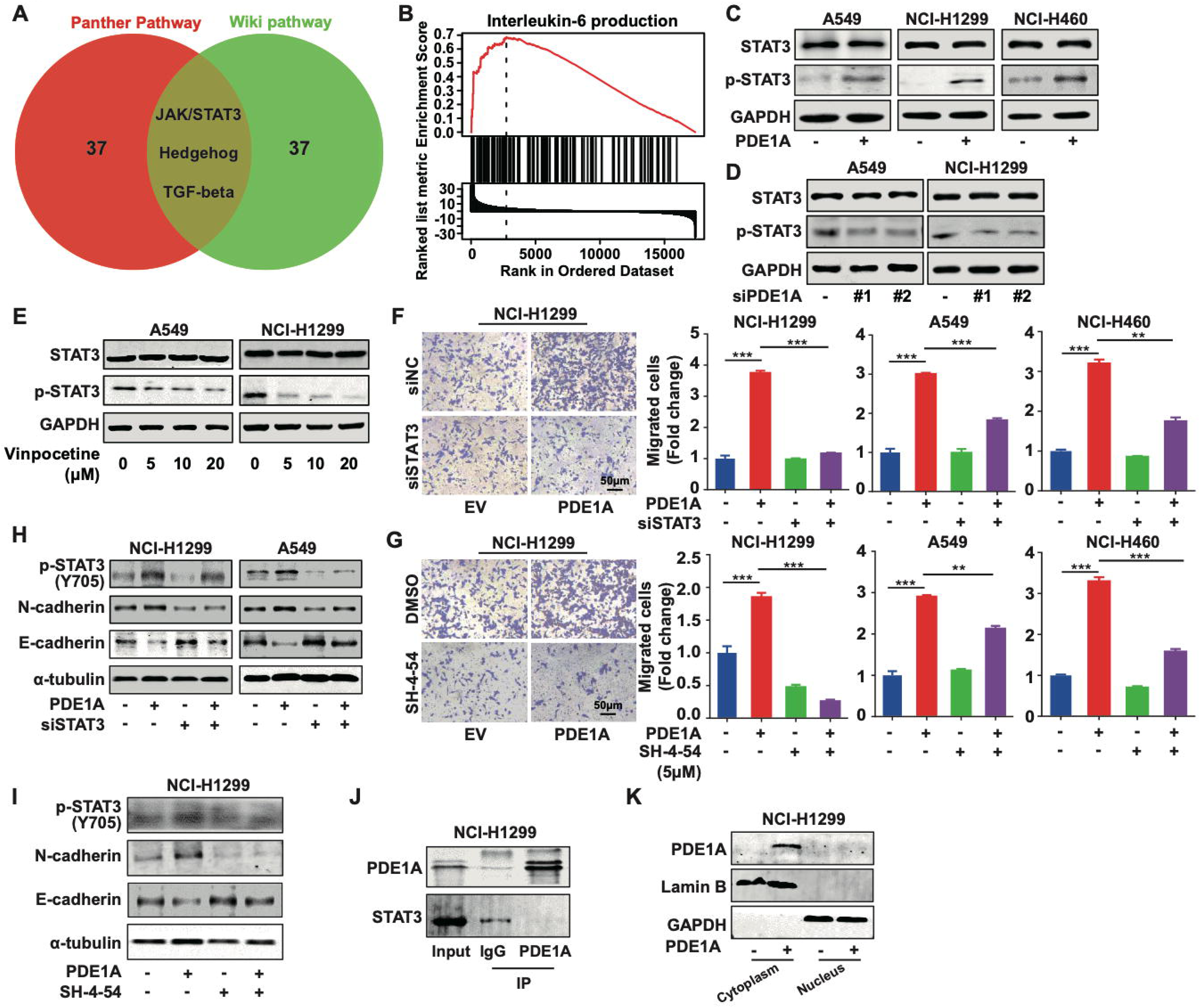
PDE1A promotes the metastasis of NSCLC cells via STAT3 signaling pathway. (A) A Venn diagram was generated using LinkedOmics, ORA was performed to analysis the molecular pathway regulated by PDE1A in NSCLC. Sample cohort: TCGA_NSCLC; Institute: UNC; Data type: RNAseq; Platform: HiSeq RNA; Attribute: PDE1A; Statistical methods: Spearman Correlation test; Patients: 515; Tools: ORA; Gene Ontology analysis: Wikipathway and Panther Pathway; Select Rank Criteria: FDR; Select Sign: Positively correlated; Significance Level: 0.05; TOP40 was selected to generate Venn diagram. (B) GSEA was performed to analyze the biological process of PDE1A in NSCLC. (C) NSCLC cells were transfected with PDE1A plasmid and empty vector for 48 h, and the expression of indicated proteins were detected. (D) NSCLC cells were transfected with control siRNA and PDE1A siRNA for 48 h, and the expression of indicated proteins were detected. (E) NSCLC cells were treated with DMSO or vinpocetine (5, 10, 20 µM) for 24 h, and the expression of indicated proteins was determined. (F) NSCLC cells overexpressing PDE1A were transfected with control siRNA and STAT3 siRNA for 48 h, the migrative abilities of NSCLC cells were determined by Transwell assay. (G) NSCLC cells overexpressing PDE1A were treated with STAT3 inhibitor SH-4-54 (5 µM) for 24 h, the migrative abilities of NSCLC cells were determined by Transwell assay. (H) NSCLC cells overexpressing PDE1A were transfected with control siRNA and STAT3 siRNA for 48 h, and the expression of indicated protein was detected by western blot. (I) NSCLC cells overexpressing PDE1A were treated with STAT3 inhibitor SH-4-54 (5 µM) for 24 h, and the expression of indicated protein was detected by western blot. (J) The interaction between PDE1A and STAT3 was determined by immunoprecipitation. (K) NCI-H1299 cells were transfected with empty vector and PDE1A overexpressing plasmid for 48 h, and the contribution of PDE1A in the cytoplasm and nucleus was determined.

Meanwhile, GSEA enrichment analysis demonstrated that PDE1A might participate in IL-6 production (Figure 5B). These findings led to the hypothesis that IL-6/JAK/STAT3 signaling may be involved in PDE1A-mediated promotion of metastasis in NSCLC.. As shown in Figure 5C, PDE1A overexpression increased the phosphorylation level of STAT3 in NSCLC cells. In contrast, PDE1A knockdown or the PDE1 inhibitor vinpocetine suppressed the phosphorylation of STAT3 in NSCLC cells (Figure 5D and 5E). Moreover, STAT3 suppression by siRNA or SH-4-54 significantly reversed the enhancement of NSCLC cell migration induced by PDE1A overexpression (Figure 5F and 5G). In addition, the suppression of STAT3 inhibited PDE1A-induced EMT progression in NSCLC cells (Figure 5H and 5I). Thus, PDE1A promoted the metastasis of NSCLC cells via activating the STAT3 signaling pathway, but the direct interaction between PDE1A and STAT3 could not be observed in NSCLC cells (Figure 5J). Moreover, PDE1A was mainly overexpressed in the cytoplasm in NSCLC cells (Figure 5K). Subsequently, the mechanism by which PDE1A promotes the STAT3 signaling pathway in the cytoplasm was further explored.

### PDE1A physically interacts with YTHDF2 and promotes the metastasis of NSCLC cells

To investigate the mechanism by which PDE1A promotes NSCLC metastasis and activates the STAT3 pathway, the proteins interacting with PDE1A in NSCLC were determined using immunoprecipitation followed by mass spectrometry analysis (Figure S10). To identify the key proteins involved in PDE1A-mediated STAT3 activation, a Venn diagram was generated, which showed that there were 9 overlapping genes among those coexpressed with STAT3 in NSCLC clinical samples, PDE1A-interacting proteins, and overexpressed in NSCLC compared with normal tissues (Figure 6A). Meanwhile, GO enrichment analysis of PDE1A interacting genes was used to predict the molecular function of PDE1A, and PDE1A might participate in m6A-containing RNA binding in NSCLC progression (Figure 6B). Based on this, it was hypothesized that PDE1A may interact with YTHDF2 and be involved in the binding of m6A-modified RNA during NSCLC progression. The physical binding between PDE1A and YTHDF2 was confirmed by silver staining and immunoprecipitation (Figure 6C and 6D). Furthermore, YTHDF2 knockdown reversed the enhancement of NSCLC migration induced by PDE1A overexpression, indicating that PDE1A might interact with YTHDF2 and promote the metastasis of NSCLC (Figure 6E and Figure S6B and S6C). The mRNA and protein levels of YTHDF2 were upregulated in NSCLC compared with normal lung tissues (Figure S7A and S7B) ^[23]^. In addition, YTHDF2 overexpression predicted poor outcomes in lung cancer patients (Figure S7C) ^[19, 20]^. Meanwhile, the activation of STAT3 by PDE1A could be reversed by YTHDF2 knockdown in NSCLC cells (Figure 6F). Furthermore, PDE1A was positively correlated with YTHDF2 expression with a Pearson correlation coefficient above 0.3 in NSCLC tissues (Figure 6G) ^[23]^. Thus, PDE1A might regulate the STAT3 signaling pathway via interacting with YTHDF2.

**Figure 6:**
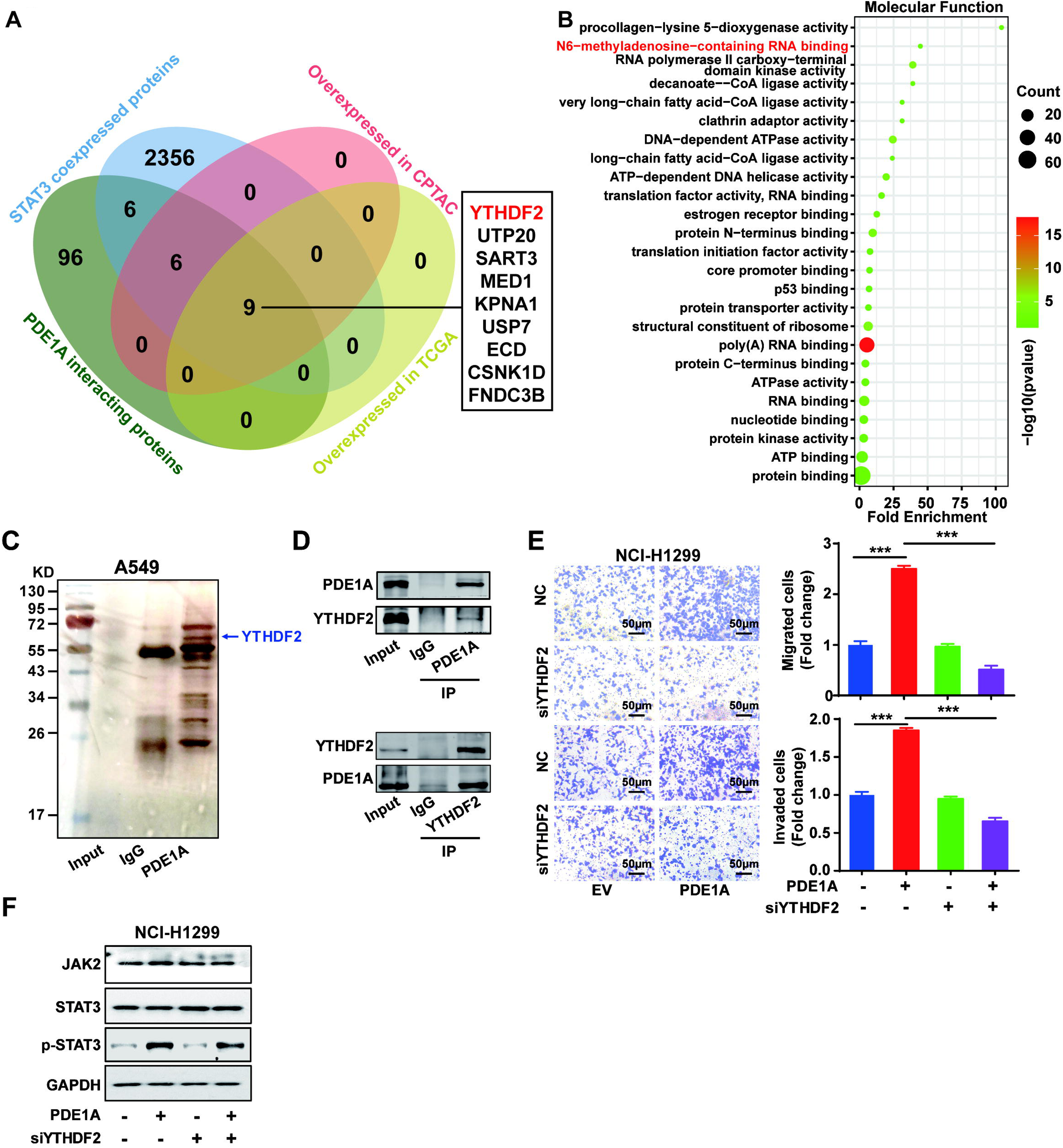
PDE1A physically interacts with YTHDF2 and promotes the metastasis of NSCLC cells. (A) Venn diagram showing the overlap among PDE1A interacting proteins (The data was collected from mass spectrometry analysis in NSCLC cells), STAT3 coexpressed genes (collected from gene correlation using UALCAN), upregulated proteins in NSCLC compared with normal tissues (analyzed by UALCAN based on CPTAC database), and upregulated genes in NSCLC compared with normal tissues (analyzed by UALCAN based on TCGA database). Pearson correlation analysis of UALCAN was used to evaluate gene correlation analyses, and Welch’s T-test was estimated to detect the significance of differences in expression levels between two groups. (B) GO enrichment analysis of PDE1A interacting genes. (C) Immunoprecipitation followed by silver staining was performed to identify protein and protein interaction using A549 cell lysate with the anti-PDE1A antibody. (D) Immunoprecipitation was used to confirm protein and protein interaction in NCI-H1299 cells. (E) NSCLC cells overexpressing PDE1A were transfected with control siRNA and YTHDF2 siRNA for 48 h, the migrative abilities of NSCLC cells were determined by Transwell assay. (F) NSCLC cells overexpressing PDE1A were transfected with control siRNA and YTHDF2 siRNA for 48 h, and the expression of indicated protein was detected by western blot.

### PDE1A interacts with YTHDF2 to regulate the SOCS2/STAT3 signaling pathway

To gain mechanistic insights into the key target mRNA through which the PDE1A/YTHDF2 complex regulates the STAT3 signaling pathway, a Venn diagram was generated, which showed that there were 33 overlapping genes among RNAs that physically interact with YTHDF2, YTHDF2-coexpressed genes, and STAT3-coexpressed genes in lung cancer (Figure 7A and 8. Figure S11) ^[24–27]^. Then, the interactions between the YTHDF2 protein and the mRNAs of 33 overlapping genes were predicted by RNA-Protein Interaction Prediction online tool. There were three predicted targets of YTHDF2 with high scores and highly correlated with STAT3 signaling as previously reported, including NRF2, SOCS2, and MET (Figure S12) ^[28]^. SOCS family members are cytokine-inducible negative regulators of the JAK/STAT pathway, and SOCS2 suppresses the binding of JAK2 and STAT3, the activity of JAK, and STAT3 activation ^[29]^. It was hypothesized that PDE1A may interact with YTHDF2, affect the stability of *SOCS2* mRNA, and thereby regulate the STAT3 signaling pathway in NSCLC cells. As shown in Figure 7B, the interaction between YTHDF2 protein and SOCS2 mRNA was confirmed by RIP, and the binding between PDE1A protein and SOCS2 mRNA was also demonstrated using RIP. Meanwhile, siPDE1A significantly enhanced the stability of SOCS2 mRNA in NSCLC cells (Figure 7C). Furthermore, YTHDF2 or PDE1A negatively regulated the expression of SOCS2 mRNA in NSCLC cells, and YTHDF2 overexpression successfully reversed the siPDE1A-induced SOCS2 mRNA accumulation. In contrast, siYTHDF2 enhanced siPDE1A-induced SOCS2 mRNA accumulation (Figure 7D). These data indicated that YTHDF2 might negatively regulate the expression of SOCS2 mRNA via cooperating with PDE1A.

**Figure 7:**
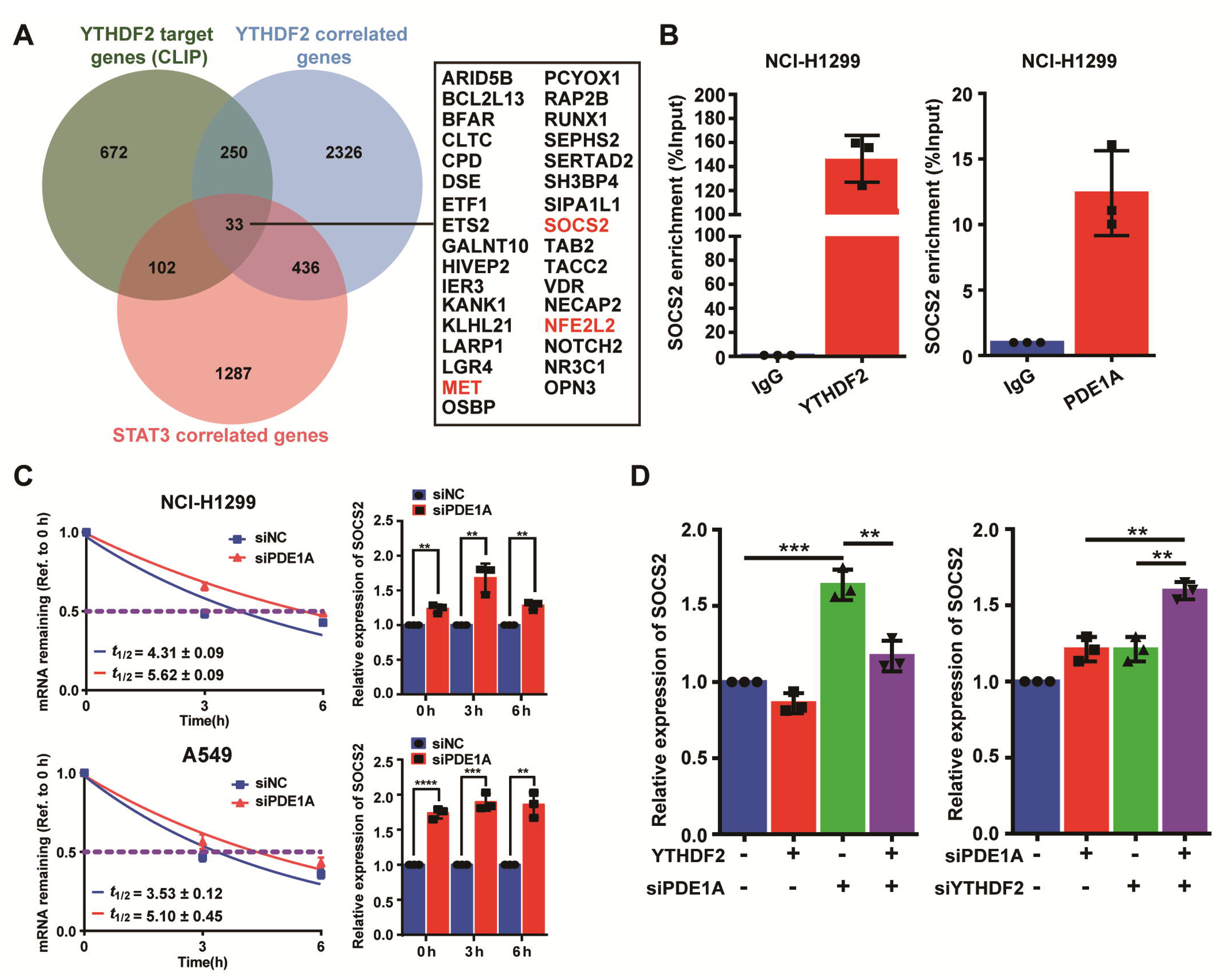
PDE1A interacts with YTHDF2 to regulate SOCS2/STAT3 signaling pathway. (A) YTHDF2-RNA complexes were identified by LC-MS/MS and collected from reference; YTHDF2 corelated genes were collected from TNMplot (https://tnmplot.com/analysis/), Gene: YTHDF2, Gene vs. all genes correlation: Genechip data, Tissue: Lung; STAT3 corelated genes were collected from cBioPortal (http://www.cbioportal.org/), Lung cancer (SMC, cancer research 2016); n=22; Gene: STAT3. The interaction between YTHDF2 protein and the mRNA of 33 overlapping genes were predicted by RNA-Protein Interaction Prediction (http://pridb.gdcb.iastate.edu/RPISeq/index.html), and the values of RF classifier and SVM classifier above 0.5 were considered positive. Comparison of the normal and the tumorous samples was performed by the Mann– Whitney U test, and Normal, tumorous and metastatic tissue gene comparison can be analyzed using Kruskal–Wallis test. (B) The interactions between protein and mRNAs were verified by RIP experiments. (C) NSCLC cells were transfected with control siRNA and siPDE1A for 48 h, and the stability of mRNA was determined by qRT-PCR. (D) NSCLC cells were transfected with control siRNA and siPDE1A for 48 h, and the expression of SOCS2 mRNA was determined by qRT-PCR.

## Discussion

NSCLC is becoming a leading cause of death globally due to its fast progression and metastatic potential, and effective therapeutic targets are urgently needed to block NSCLC metastasis ^[30]^. PDEs are regarded as therapeutic targets for multiple diseases, but the feasibility of targeting PDEs to treat NSCLC metastasis may need further investigation ^[31]^.The present investigation provides the first evidence that PDE1 promotes metastasis and epithelial-mesenchymal transition (EMT) progression in NSCLC cells. Furthermore, the expression of PDE1A was closely correlated with the disease progression of NSCLC. These data first indicated with PDE1A might be an efficacious therapeutic target for patients with metastatic NSCLC.

The stimulation of the PDEs requires physiological concentrations of Ca^2+^ and calmodulin, and Ca^2+^/calmodulin-dependent cyclic nucleotide PDE1 is involved in the communication between the cyclic nucleotide and Ca^2+^ second messenger systems ^[32]^. The increasing of Ca^2+^ in cancer cells stimulates exosome biogenesis and release under both physiological and pathological conditions ^[33, 34]^. In the context, it was hypothesized that PDE1A may be involved in exosome biogenesis and release in NSCLC cells, playing a crucial role in intercellular communication within the tumor microenvironment (TME). GSEA showed that PDE1A might be involved in angiogenesis, vasculature development, and blood vessel development. Indeed, NSCLC cells overexpressing PDE1A promoted angiogenesis in the TME, and PDE1A knockdown significantly suppressed angiogenesis of NSCLC *in vivo*. Furthermore, an exosome release inhibitor successfully reversed the angiogenesis promoted by NSCLC cells overexpressing PDE1A. Thus, the present study first illustrated that PDEs might play an important role in angiogenesis and the TME via regulating exosome biogenesis and release in cancer cells. Analysis of PDE1A co-expressed genes in NSCLC revealed a significant enrichment of the IL-6/JAK/STAT3 signaling pathway, suggesting its involvement as a downstream pathway of PDE1A. Furthermore, PDE1A promoted the metastasis of NSCLC cells via the STAT3 signaling pathway. Targeting the IL-6/JAK/STAT3 signaling pathway is considered as a promising therapeutic strategy for the management of NSCLC ^[35]^. However, the direct interaction between PDE1A and STAT3 could not be observed in NSCLC cells. Subsequently, the mechanism by which PDE1A promotes the STAT3 signaling pathway was investigated. It demonstrated that PDE1A interacts with YTHDF2 and contributes to NSCLC progression, with the interaction between YTHDF2 and PDE1A being verified for the first time in NSCLC cells. Meanwhile, YTHDF2 might act as an m6A RNA “reader” by interacting with PDE1A, but the mechanism might need further investigation. YTHDF2 destabilizes mRNAs via degrading target transcripts, but it also stabilizes important oncogenic drivers, such as *MYC* and *VEGFA* transcripts, in an m6A-dependent manner ^[36]^. It was demonstrated that YTHDF2 destabilized SOCS2 mRNA via interacting with PDE1A, but the mechanism by which YTHDF2 sorts mRNA might need further investigation. In addition, it is worth testing if PDE1A inhibition affects metastasis in lung cancer models and sensitizes cisplatin in resistant NSCLC cells in vitro and in vivo. The role of YTHDF2 in PDE1A-driven tumor metastasis should be elucidated in future studies.

Collectively, the results suggest that PDE1A promotes metastasis in NSCLC cells. Furthermore, PDE1A overexpression is correlated with angiogenesis and poor outcomes of NSCLC patients. In addition, PDE1A interacts with YTHDF2 and regulates the JAK/STAT3 signaling pathway via degrading SOCS2 mRNA. Therefore,it reveals the effect and mechanism of PDE1A in promoting NSCLC metastasis, the data not only uncovers a novel PDE1A/YTHDF2/STAT3 signaling pathway in NSCLC progression but also provides novel therapeutic strategies to treat NSCLC patients with metastasis.

## Materials and methods

### Materials

Vinpocetine (V107535) was purchased from Aladdin (Shanghai, China). SH-4-54 (S7337), SQ22536 (S8283), and GW4869 (S7609) were obtained from Selleck Chemicals (Houston, TX, USA). Antibodies against β-actin (SC-1616) and GAPDH (SC-25778) were obtained from Santa Cruz Biotechnology (Dallas, Texas, USA). Antibodies against N-cadherin (14215), STAT3 (9139), p-STAT3 (Y-705) (9145), Vimentin (5741), and JAK2 (3230S) were obtained from Cell Signaling Technology (Danvers, MA, USA). An antibody against α-tubulin (AT7819) was purchased from Beyotime Biotechnology (Shanghai, China). Antibodies against PDE1A (12442-2-AP), YTHDF2 (24744-1-AP), lamin B (12595-1-AP), and E-cadherin (20874-1-AP) were obtained from Proteintech (Rosemont, IL, USA).

### Cell lines and cell culture

Human NSCLC cell lines (A549, NCI-H1299, and NCI-H460), lung fibroblasts (HELF), and umbilical vein endothelial cells (HUVECs) were maintained in RPMI-1640 medium supplemented with 10% fetal bovine serum (FBS) and 100 U/ml penicillin/streptomycin. All the cell lines were purchased from the Shanghai Institute of Biochemistry and Cell Biology (Shanghai, China), cultured at 37 °C with 5% CO_2_ and confirmed to be mycoplasma-free.

### siRNA transfection

Scramble siRNA, siPDE1A, siPDE1B, siPDE1C, siSTAT3, and siYTHDF2 were synthesized by GenePharma (Suzhou, China). Then, NSCLC cells were transfected with siRNA (40 nM) using PolyPlus-transfection reagent in accordance with the instructions. The sequences of siRNAs are summarized in Figure S8.

### Sulforhodamine B (SRB) assay

NSCLC cells were treated with the indicated compounds, and subsequently fixed with ice-cold TCA and stained with 0.4% SRB (w/v) solution. Cell proliferation was determined by SRB assay according to previously reported methods ^[14]^.

### Wound healing assay

NSCLC cells were seeded on a 24-well plate and cultured as a monolayer to 90% confluence. The monolayer was scratched with a 10 µL pipette tip, and then the cells were cultured with FBS-free culture medium for 24 h. Images of the wounded cell monolayer were taken using a microscope (Olympus, Japan) at 0 h and 24 h. The wound closure rate was calculated as follows: (G_0_-G_24_) / G_0_ × 100%, where G_0_ and G_24_ represent the gap areas at 0 and 24 h, respectively.

### Migration and invasion assays using transwells

Migration and invasion assays were performed using a 24-well transwell chamber system (pore size: 8 µm, Corning, USA). A total of 5 × 10^4^ cells were seeded in the upper chamber of an insert with 0.4 ml serum-free culture media in 24-well plates. Then, 0.6 ml culture medium with 20% FBS was added to the lower chamber. For invasion assays, the upper transwell chamber of the insert was coated with Matrigel (BD Biosciences, Bedford, MA, USA) before plating cells. 50 μl of Matrigel was dissolved in 450 μl of culture medium and added 100 μl solution into the upper transwell chamber. After incubation for 24 h, migratory or invasive cells were stained with 0.5% crystal violet and analyzed under a light microscope.

### Protein extraction and western blotting analysis

Briefly, cells were washed three times with cold PBS and pelleted. The pellet was resuspended in lysis buffer (NP40 lysate), incubated on ice with frequent vortex for 30 min, and the lysate was obtained by centrifugation at 10000 × g for 30 min. Proteins were fractionated by SDS-PAGE, transferred onto PVDF membranes, blocked in 5% nonfat milk in PBS/Tween-20, and then blotted with specific primary antibody overnight at 4 °C, followed by incubation with secondary antibody for 1□h at room temperature. Bands were detected by the Odyssey CLX Image Studio system (version 5.0.21, LiCor Odyssey, LI-COR Biosciences).

### Immunoprecipitation and LC-MS/MS

For coimmunoprecipitation, the cells were lysed with IP buffer (20 mM HEPES, 25% glycerine, 210 mM NaCl, 1.5 mM MgCl_2_, 0.05 mM EDTA, 0.2% NP40, 1 × cocktail, 1 mM PMSF, 2 mM DTT) and centrifuged at 10000 × g for 30 min at 4 °C. The cell lysates were treated with protein G magnetic beads at 4 °C for 1 h. Subsequent immunoprecipitation reactions were set up with equal quantities of the lysates. Primary antibody was added to the lysate, and the mixture was incubated overnight with slow shaking at 4 °C, and then incubated with protein G magnetic beads at 4 °C for 1 h. Subsequently, the lysates were centrifuged at 3000 g at 4 °C for 5 min. The supernatant was aspirated, the protein G magnetic beads were washed 3–4 times with lysis buffer, and detection was performed using SDS-PAGE or LC-MS/MS (Micrometer Biotech Company, Hangzhou, China).

### Extraction of RNA and quantitative real-time PCR (qRT-PCR)

Total RNA was extracted from cells by utilizing TRIzol™ Reagent (Invitrogen, Thermo Fisher Scientific). RNA was reverse-transcribed into cDNA using an iScript cDNA Synthesis kit (Bio-Rad, Hercules, CA, USA). qRT-PCR was performed using the PerfectStart Green qPCR SuperMix kit (TransGen Biotech, Beijing, China). The levels of mRNA were analyzed by the Bio-Rad CFX96 real-time PCR system (Bio-Rad, Hercules, CA, USA). GAPDH was used as an endogenous control for mRNA qualification and the 2-ΔΔCt method was applied to calculate the relative expression. The primers used are listed in Figure S9.

### RNA binding protein immunoprecipitation (RIP)

RIP assays were performed using a Magna RIP Kit (17-701, Millipore, MA, USA) following the manufacturer’s instructions. In brief, magnetic beads precoated with 5□μg normal antibodies against PDE1A/YTHDF2 or rabbit IgG (Millipore) were incubated with cell lysates at 4□°C overnight. The beads containing immunoprecipitated RNA-protein complexs were treated with proteinase K to remove proteins. Then RNAs of interest were purified with TRIzol and measured by qRT-PCR.

### Plasmid and shRNA transfection and infection

Two micrograms of overexpressing plasmid or shRNA of the indicated genes was transfected into cells using PolyPlus-transfection reagent. For shRNA used in lentivirus-mediated interference, complementary sense and antisense oligonucleotides encoding shRNAs targeting PDE1A were synthesized, annealed, and cloned into pLKO.TRC vector (Addgene, 10878). The PDE1A-FLAG overexpression plasmid was synthesized by GenScript (Nanjing, Jiangsu, China). YTHDF2-HA expression plasmids were synthesized via cloning YTHDF2 with an HA tag into the pcDNA3.1(−) vector.

### *In vivo* animal experiment

Female nude mice (BALB/c, 4-6 weeks old) were obtained from Shanghai SLAC Laboratory Animal Co., Ltd. A total of 2□×□10^6^ NCI-H1299 cells transfected with shPDE1A/control shRNA or PDE1A overexpressing plasmid/empty vector were suspended in 0.1□ml of PBS and injected into mice via the tail vein. After 60 days, the mice were sacrificed, and the lung tissues were collected to observe pulmonary nodules. The lung, liver, kidney, pancreas, and other tissues were separated, fixed in 4% paraformaldehyde, and stained using H&E. All animal procedures were conducted in accordance with the guidelines and regulations approved by the Institutional Animal Care and Use Committee (IACUC) of Zhejiang University City College. Ethical approval for the study was obtained under protocol number 22001.

### mRNA stability assay

NSCLC cells were seeded in six-well plates and grown to approximately 30% confluence followed by siRNA transfection and incubation for 24 h. Then, cells were incubated with actinomycin D (5□μg/ml) for 0, 3 or 6 h followed by RNA extraction. The half-life of mRNA was analyzed by qRT-PCR. The mRNA expression for each group at the indicated time was calculated and normalized to GAPDH.

### Statistical analysis

Data are presented as the mean□±□SD from 3 independent experiments. Two-tailed Student’s t test was used to compare 2 groups. P values < 0.05 were considered significant. *P < 0.05; **P < 0.01; ***P < 0.001.

## Acknowledgements

The study was funded by Huadong Medicine Joint Funds of the Zhejiang Provincial Natural Science Foundation of China (LHDMY22H160001), Natural Science Foundation of Ningbo City (2022J206), National Natural Science Foundation of China (82273352), the Youth Incubation Project of Xian Health Commission (2024qn01), Scientific Research Foundation of Hangzhou City College (X-202305), National College Student Innovation and Entrepreneurship Project (202313021036) and Zhejiang Provincial Medical and Health Technology Project (2025KY391).

## Conflict of Interest

The authors have declared that no competing interest exists.

## Availability of data and material

The data presented in the study are included in the article and additional material.

## Ethics approval and consent to participate

The experiments were performed in compliance with the National Institutes of Health Guide for the Care and Use of Laboratory Animals.

## Consent for publication

Not applicable.

## Supplementary information

1. Supplementary file 1: Figure S1. tif

The expression of PDE1A in normal lung (A) and lung cancer (B). The data were obtained from The Human Protein Atlas (https://www.proteinatlas.org).

2. Supplementary file 2: Figure S2. tif

The biological processes related to PDE1A in NSCLC is predicted using the LinkedOmics. Data collected from LinkedOmics (www.linkedomics.org/admin.php) are shown. ORA (A) and GSEA (B) were performed to analyze the biological processes related to PDE1A in NSCLC. Sample cohort: TCGA_NSCLC; Institute: UNC; Data type: RNAseq; Platform: HiSeq RNA; Attribute: PDE1A; Statistical methods: Spearman Correlation test; Patients: 515; Tools: ORA and GSEA; Gene Ontology analysis: biological process. (C) GSEA demonstrated that PDE1A participated in mesenchyme development and the cellular response to VEGF stimulus.

3. Supplementary file 3: Figure S3. tif

The expression of PDE1A does not affect the proliferation of NSCLC cells. (A-B) NSCLC cells were transfected with PDE1A siRNA or control siRNA (A) and empty vector or PDE1A plasmid (B) for 24 h in 6-well plates, then transferred to 96-well plated for the indicated times, and finally subjected to SRB assay.

4. Supplementary file 4: Figure S4.tif

PDE1A silence suppresses the metastasis of NSCLC cells. (A) NSCLC cells were transfected with control siRNA, siPDE1A, siPDE1B, and siPDE1C for 24 h, and cells were transferred to Transwell chambers without a Matrigel coating on the insert membrane, and the cell migration were determined after 24 h. (B) NSCLC cells were transfected with indicated siRNA for 48 h, and the expression of the indicated proteins were detected. (C) The lungs of nude mice were separated and stained with Bouin’s solution and the pulmonary metastatic nodules were counted. (D) The relative body weight of each group in the in vivo metastasis studies in figure 2I.

5. Supplementary file 5: Figure S5.tif

PDE1A promotes the metastasis of NSCLC cells. (A) The migrative abilities of P3 and P0 NSCLC cells were determined. (B) The mRNA levels of indicated genes were determined in P3 and P0 NSCLC cells using qRT-PCR. (C) Before the NSCLC cells were injected into the nude mice, the migrative abilities of NSCLC cells were determined using Transwell assay. (D)The relative body weight of each group in the in vivo metastasis studies in figure 3H.

6. Supplementary file 6: Figure S6. tif

PDE1A promotes the metastatic potential in a cAMP-independent manner. (A) NSCLC cells were transfected with control siRNA and PDE1A siRNA, after 24 h cells were collected for Transwell assay, cells were treated with DMSO or 5 µM SQ22536 for 24 h, and the migrative abilities of NSCLC cells were determined. (B-C) NSCLC cells were transfected with control siRNA and YTHDF2 siRNA for 48 h, and the knockdown efficiency of YTHDF2 was confirmed by western blot (B) and qRT-PCR (C).

7. Supplementary file 7: Figure S7. tif

Overexpression of YTHDF2 predicts poor outcomes for lung cancer patients. (A-B) The mRNA and protein levels of PDE1A in NSCLC and normal lung tissues are shown. Data collected from UALCAN was shown. Gene: YTHDF2; TCGA dataset: Lung adenocarcinoma (A); CPTAC dataset: Lung adenocarcinoma (B). (C) The prognostics value of YTHDF2 in NSCLC patients was identified using PROGgeneV2 online tool (www.progtools.net/gene).

8. Supplementary file 8: Figure S8

The siRNA sequence used for knocking down the indicated genes.

9. Supplementary file 9: Figure S9

Primers sequences for detecting the expression of the indicated genes.

10. Supplementary file 10: Figure S10

Identification of PDE1A protein interactions by mass spectrometry.

11. Supplementary file 11: Figure S11

The potential target genes of YTHDF2 in NSCLC cells.

1. sample Supplementary file 12: Figure S12

The potential target genes of YTHDF2 in NSCLC cells.

## Notes

### Competing Interest Statement

The authors have declared no competing interest.

### Summary of Updates

Impact statements were revised; the supplementary files were renamed; the author contributions were removed

