## Supplementary figures and images for "Phosphodiesterase 1A physically interacts with YTHDF2 and reinforces the progression of non-small cell lung cancer"

### Supplementary Figure 1

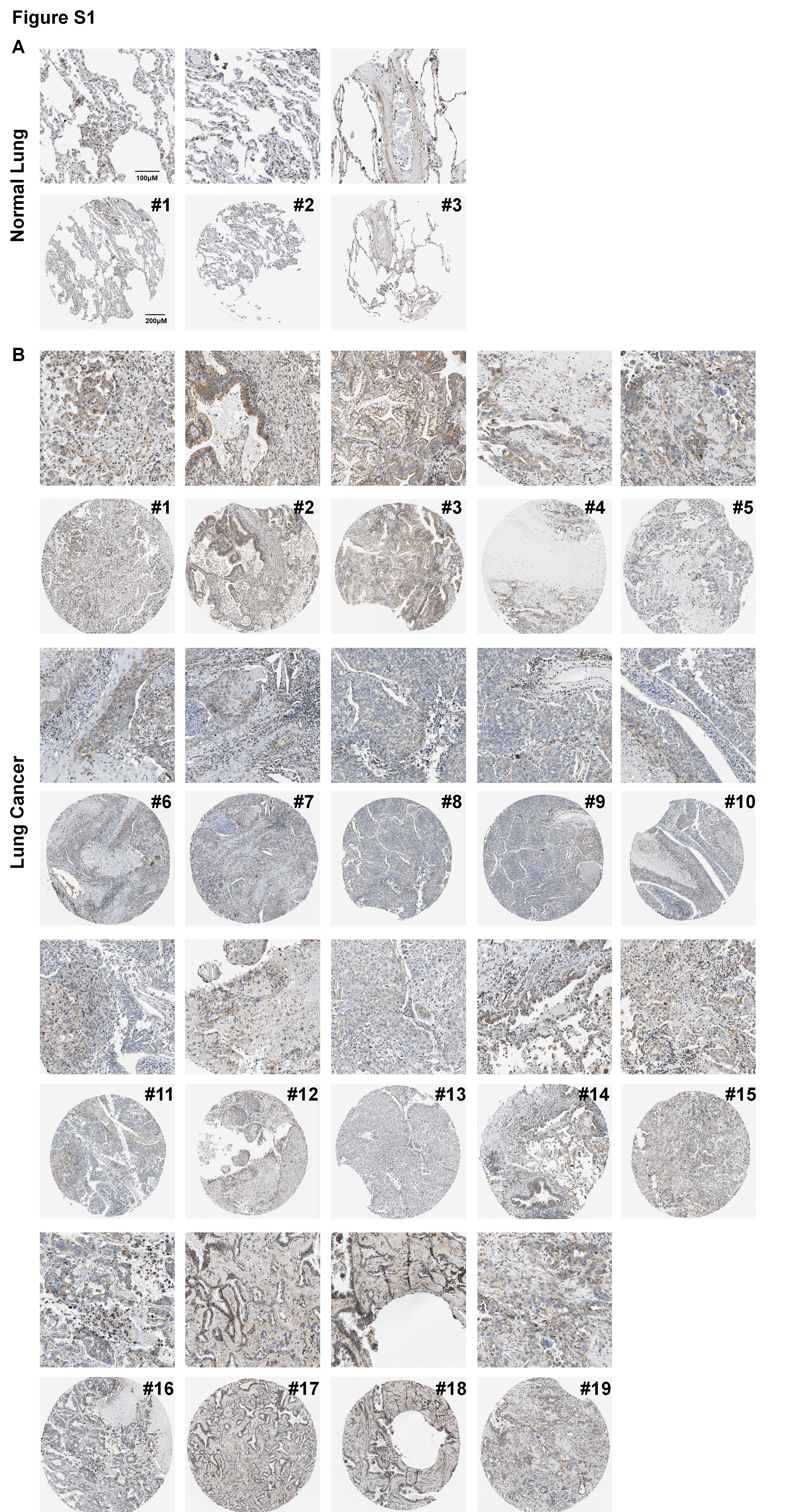

### Supplementary Figure 2

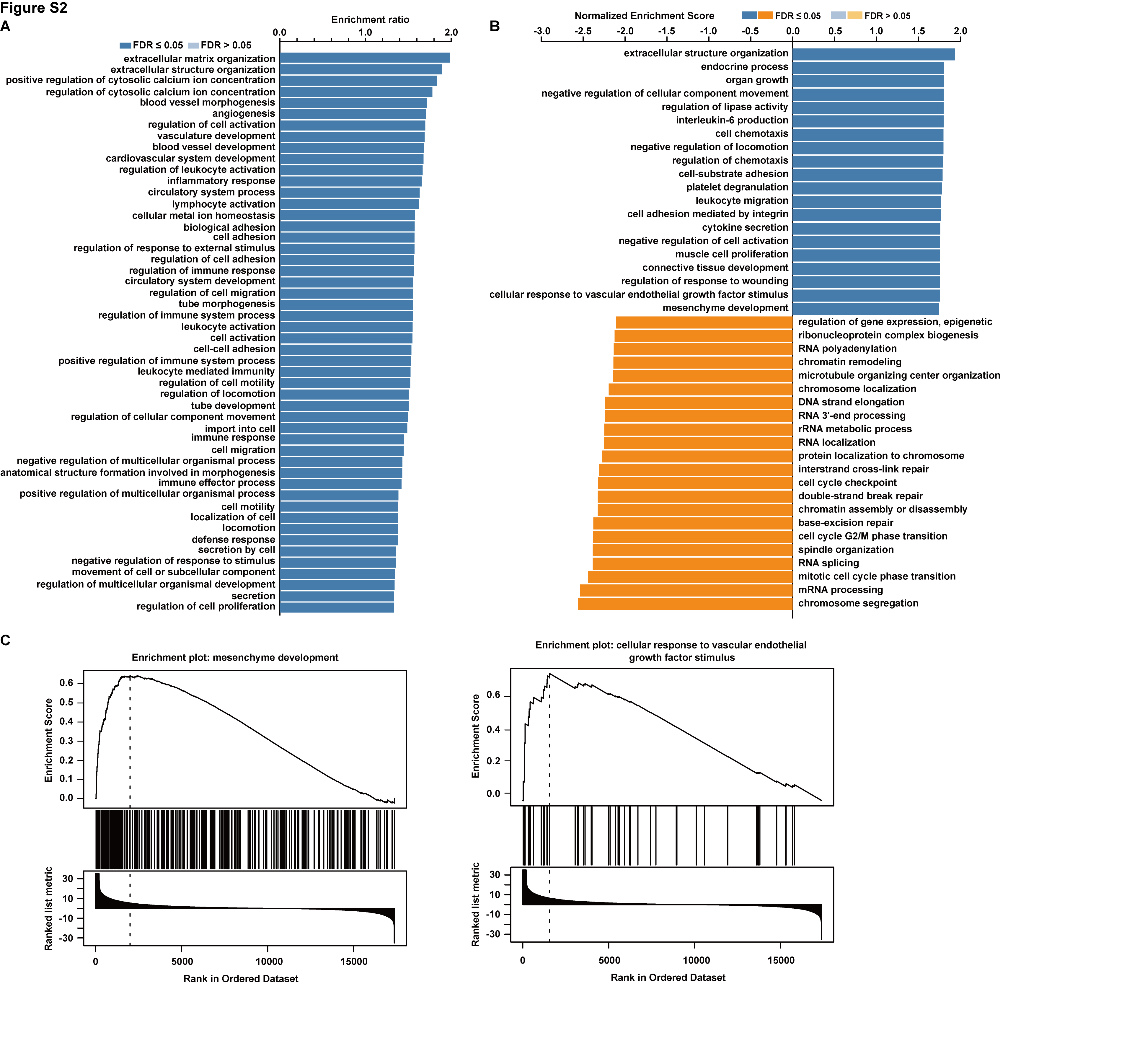

### Supplementary Figure 3

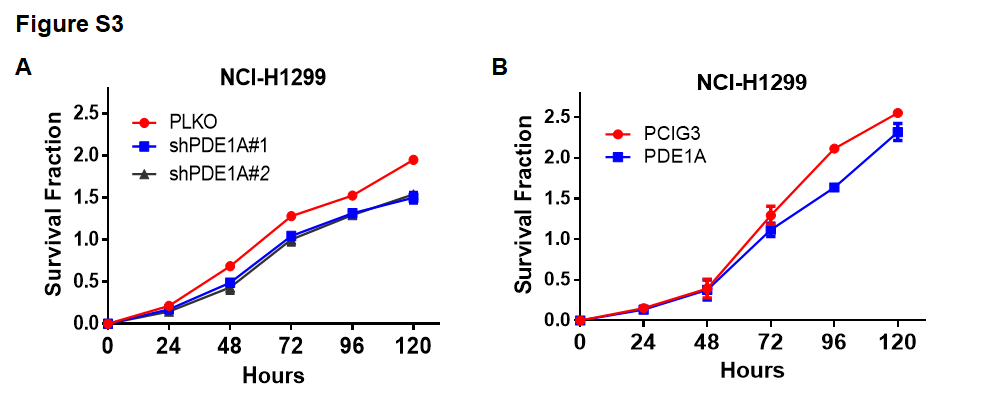

### Supplementary Figure 4

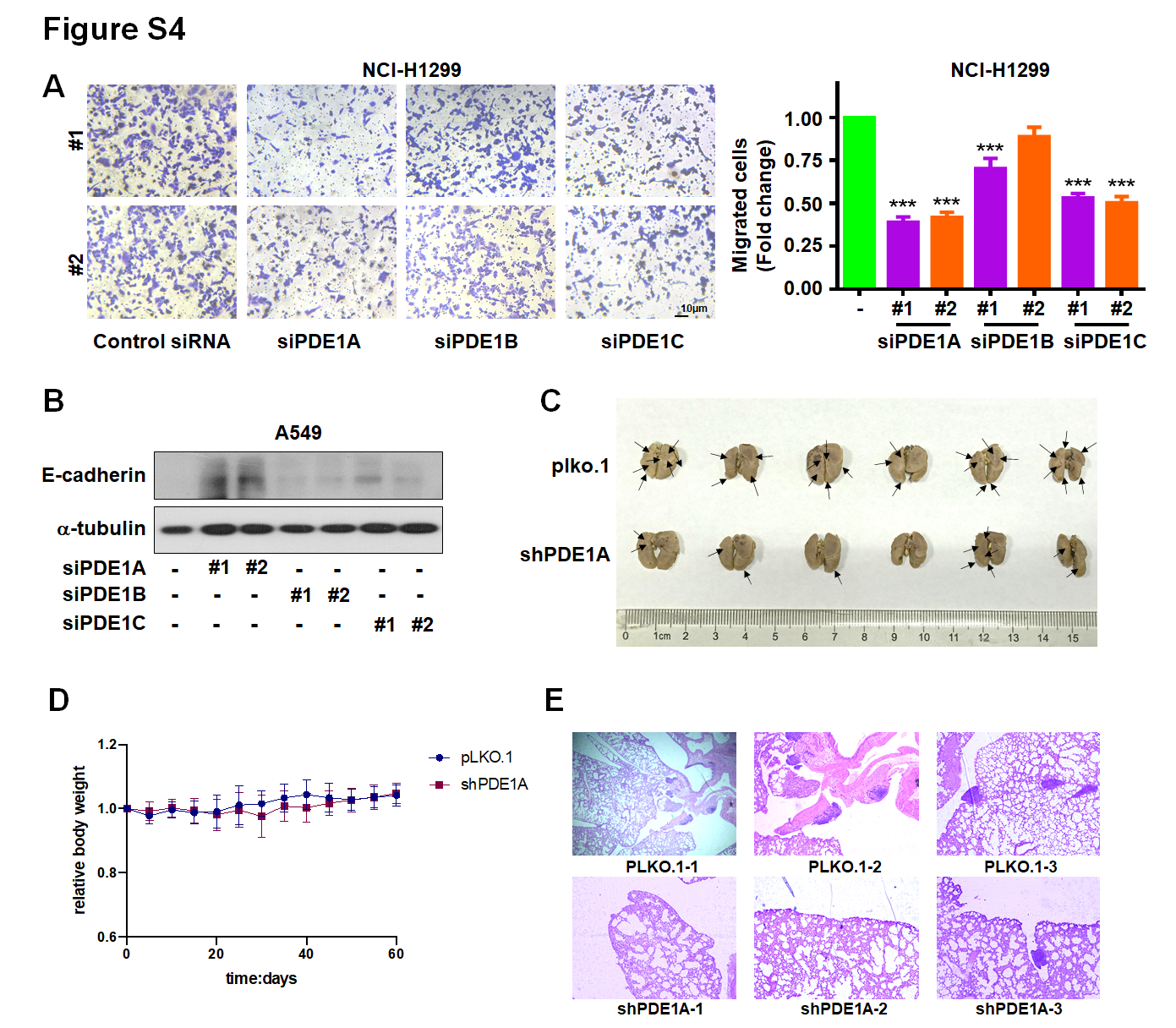

### Supplementary Figure 5

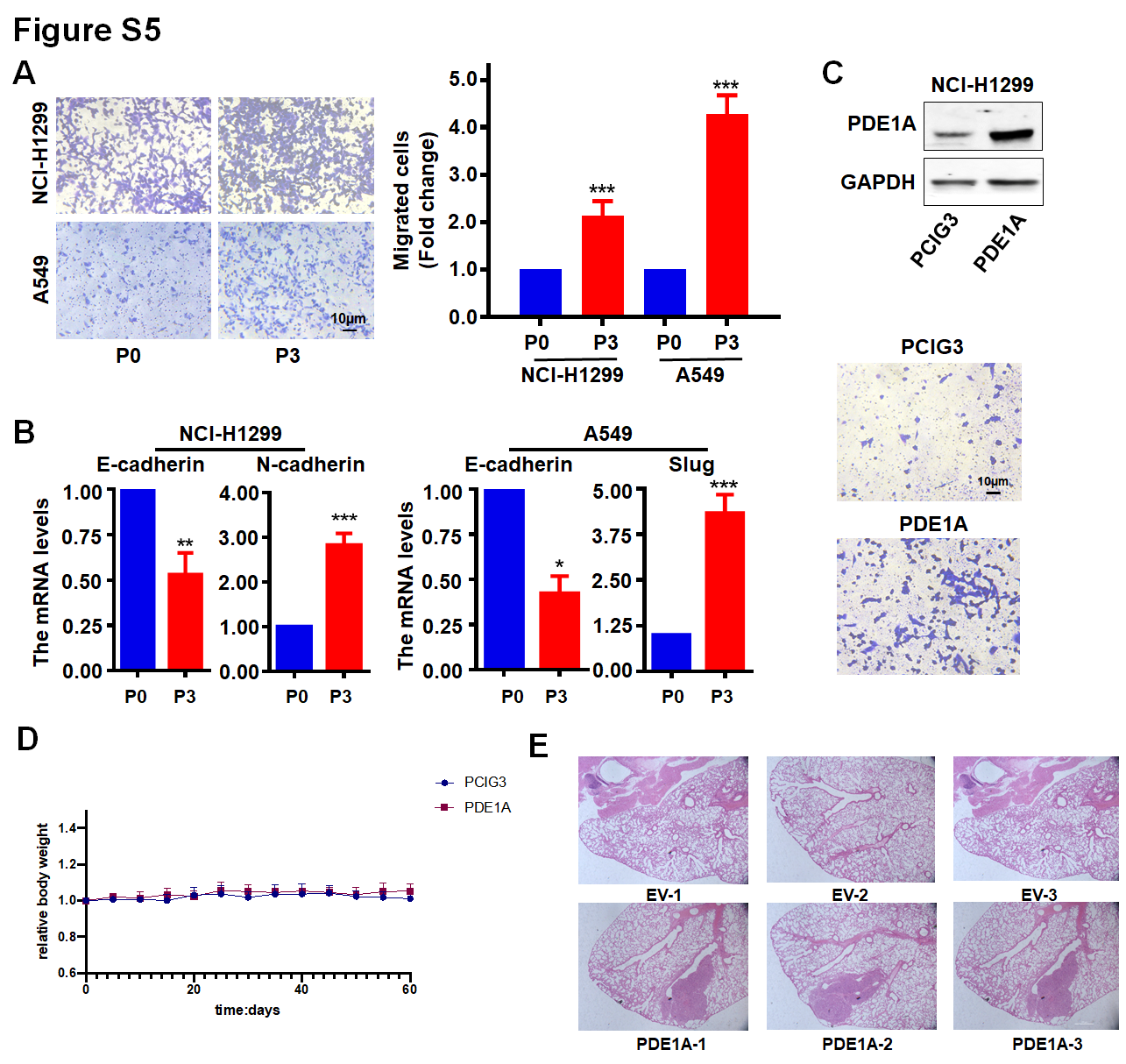

### Supplementary Figure 6

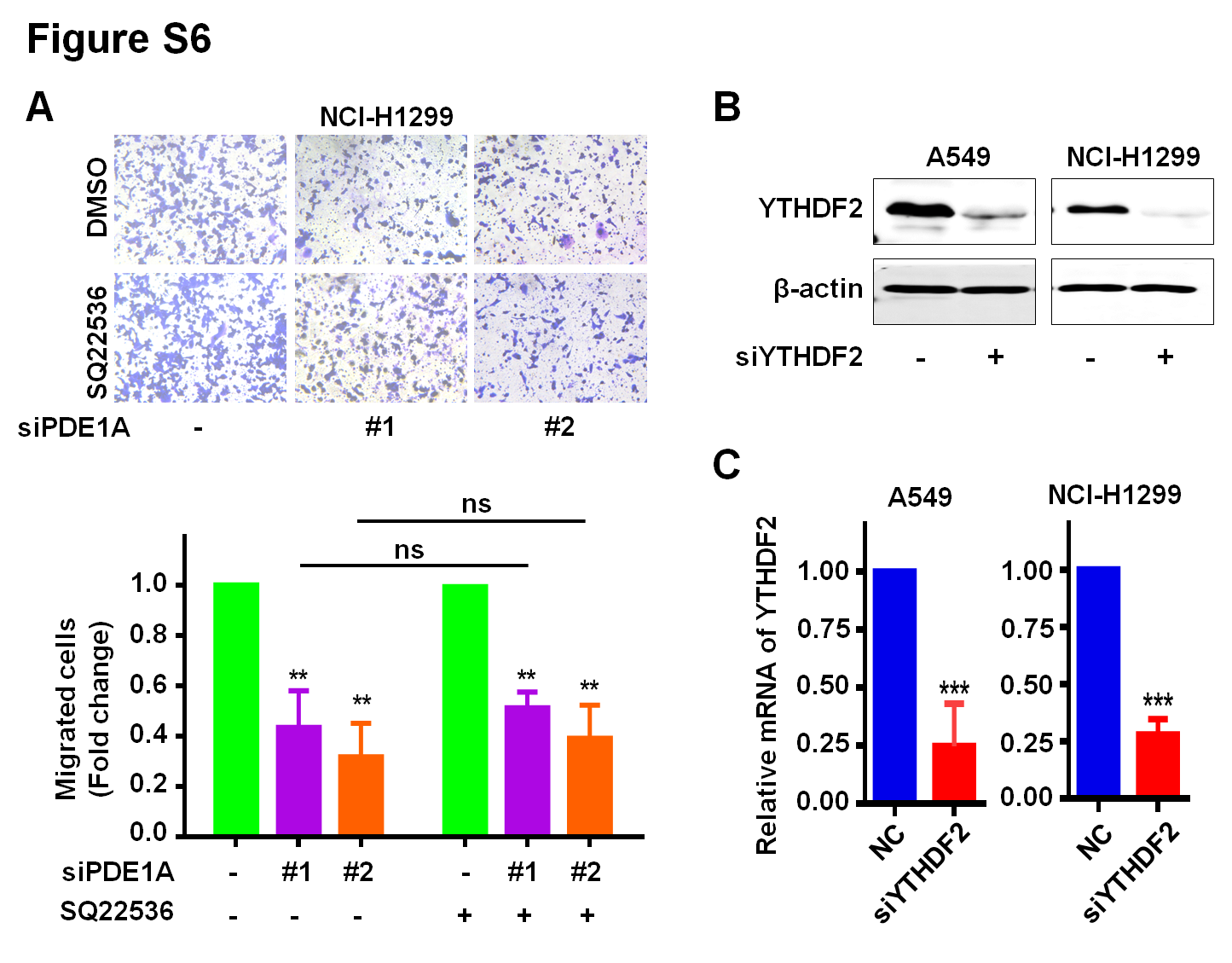

### Supplementary Figure 7

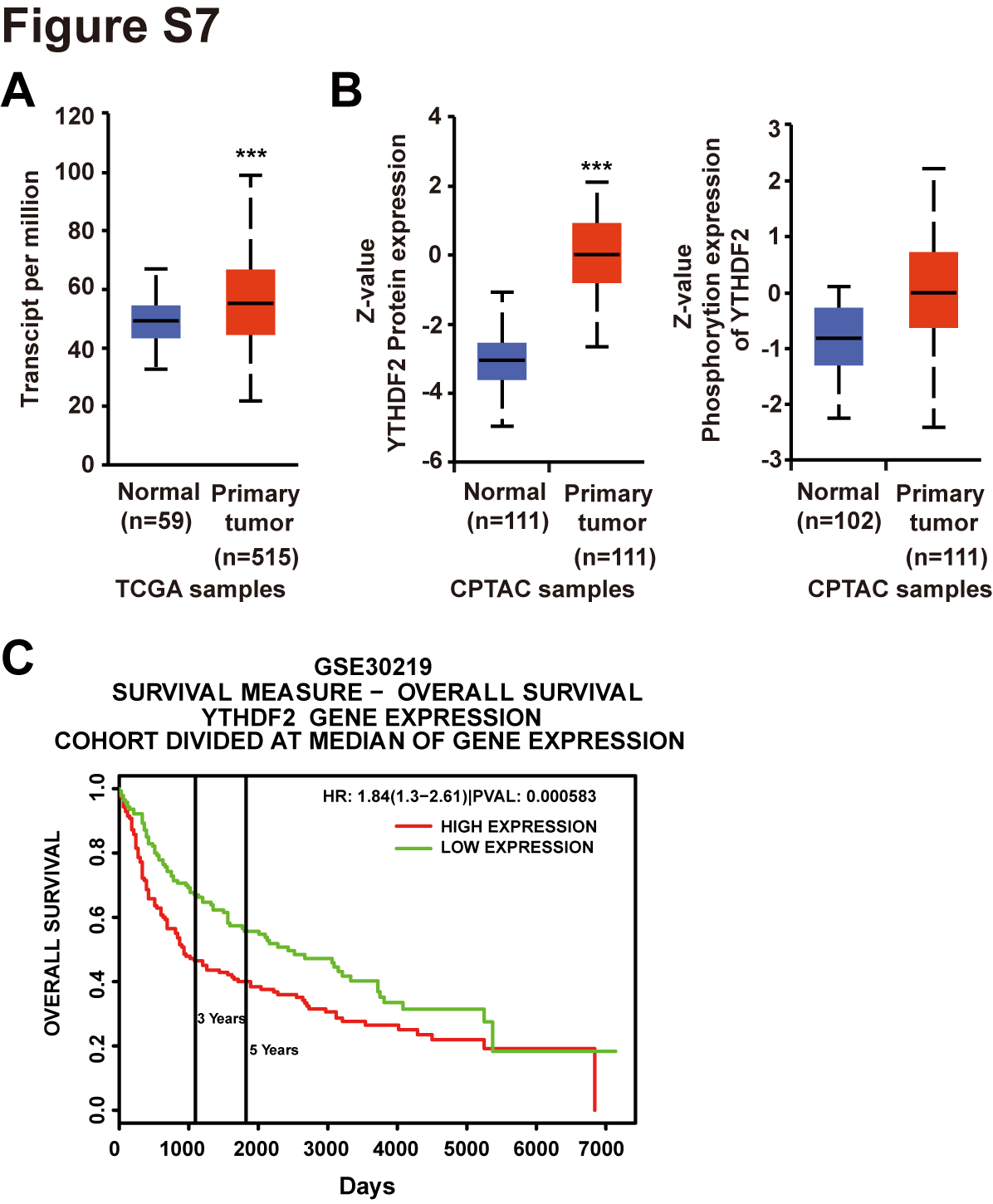
